# Marbled Murrelet Habitat Suitability in Redwood Timberlands of Northern Coastal California Utilizing LiDAR-Derived Individual Tree Metrics

**DOI:** 10.1101/2024.08.13.607789

**Authors:** David W. Lamphear, Keith A. Hamm, Desiree A. Early, Trent L. McDonald

## Abstract

The marbled murrelet (*Brachyramphus marmoratus*) is a threatened seabird found from Southern California to Alaska that forages at sea but nests in near coastal forests. Marbled murrelet nesting habitat is generally comprised of old-growth or mature forests with large trees having platforms suitable for nesting. Here, we estimate a habitat suitability model (HSM) that relates evidence of nesting to characteristics of putative trees derived from high resolution light imaging detection and ranging (LiDAR) data. Our study area in Northern California contained stands of old-growth forests on state, federal, and private lands but was predominated by private second-growth redwood and Douglas-fir timberlands. We estimated a two-sample HSM using Maxent software and implemented objective and repeatable covariate selection, model evaluation, and classification methods. Our HSM predicts relative likelihood of occupancy using predicted Habitat Suitability Index (HSI) values that we then classify into five habitat classes based on a novel use the of predicted to expected (P/E) ratio curve. From HSI predictions, we identified patches of murrelet habitat and estimated concave polygons surrounding individual trees within and proximately close to each patch. These methods provide repeatable boundaries for identification of patches and important individual trees based on HSI. Patches with greater relative probability of occupancy were characterized by high densities of 60-meter and taller trees, a large sum of heights for 50-meter and taller trees, and high values of standard deviation of 50-meter and taller trees. During hold-out model evaluations, our HSM showed extremely high fidelity for known patches with indirect evidence of nesting based on occupancy.

## Introduction

The United States Fish and Wildlife Service listed the marbled murrelet (*Brachyramphus marmoratus*) as threatened under the federal Endangered Species Act in 1992. That same year, the state of California listed the bird as endangered under the California Endangered Species Act. At the time, scientists knew that this seabird foraged offshore and nested in nearshore older forests along the Pacific Coast of the United States and Canada. Since then, several studies have investigated landscape and stand-level characteristics that comprise suitable marbled murrelet nesting habitat. Several studies of landscape scale habitat use found associations between marbled murrelet nesting and large amounts of unfragmented old-growth or mature forests (McShane et al., 2004; Raphael et al., 2016a; Oregon Department of Fish and Wildlife, 2018). Other studies characterized marbled murrelet nesting habitat as un-fragmented old-growth low elevation sites within the fog zone near other marbled murrelet nesting sites (Hamer et al., 1994; Meyer et al., 2002, 2003; Raphael et al., 2016b; Valente et al., 2021). Still other studies found a negative association between marbled murrelet nest sites and increasing amounts of forest fragmentation, edge, and areas of logged and immature forests (Burger, 2001; Burger, 2002; Cullen, 2002; McShane et al., 2004; Nelson et al., 2006; Baker et al., 2006; Raphael et al., 2016b).

At a smaller scale, individual forest stands containing marbled murrelet nests tend to have high densities of large trees (minimum diameter at breast height of 120 centimeter (4 feet)) with multiple platforms, multiple canopy layers, canopy gaps, high epiphyte thickness, high epiphyte percentages, and greater tree heights (Hamer, 1995; Hamer and Nelson, 1995; Grenier and Nelson, 1995; Miller and Ralph, 1995; Hamer and Meekins, 1996; Perez-Comas and Skalski, 1996; Nelson and Wilson, 2002; Nelson et al., 2002; Meyer et al., 2003; Baker et al., 2006). In one Oregon study, observed nest trees contained 8-92 platforms (Oregon Department of Fish and Wildlife 2018). Other Oregon studies documented few nests in non-old growth trees (60-80 years) with dwarf mistletoe, deformations, or structural elements that support a nest (Nelson and Wilson, 2002; Oregon Department of Fish and Wildlife, 2021). In addition, canopy complexity is an important component of nesting stands because openings are required for landing and takeoff (Burger, 2002; Nelson et al., 2006; Hamer et al., 2021). In California, marbled murrelets nest primarily in old growth redwood and Douglas-fir forests (Miller and Ralph, 1995; Meyer et al., 2002; Meyer et al., 2004; Baker et al., 2006; Hébert and Golightly, 2006; Golightly et al., 2009). Carter and Erickson (1988) documented marbled murrelets nesting up to 40 kilometers inland from the northern California coast. This literature provides a basis for identifying potential marbled murrelet habitat using stand-level criteria, but we wanted to develop a reliable tool for identifying a continuum of probable marbled murrelet habitat across a landscape at scales smaller than the typical stand.

In our study, we had three objectives. First, as part of a federal habitat conservation plan (HCP; https://www.fws.gov/endangered/what-we-do/hcp-overview.html), we developed a correlative habitat suitability model (HSM) for marbled murrelets to identify, at the patch and study area scale, forested areas that correlate well with areas where marbled murrelets have exhibited indirect evidence of nesting (“occupied behaviors”; Evans Mack et al., 2003) or where other observations provided direct evidence of nesting. Second, we developed habitat classes based on the predicted to expected (P/E) ratio curve of HSI values (Boyce et al., 2002; Hirzel et al., 2006). Third, we estimated concave polygons surrounding the patch and individual trees that influence nesting potential of an area and used these polygons to delineate management-friendly patch boundaries and associated habitat quality predictions. These patch boundaries would include trees in the core area of nesting patches but also included peripheral large trees that contributed to nesting suitability. To our knowledge, this would be the first successful use of LiDAR derived height and density of individual trees to model marbled murrelet nesting habitat.

### Study Area

Our study area, comprised of Green Diamond Resource Company lands plus a ½ mile buffer, was located on a mix of commercial timberlands, state, federal, and Tribal lands, with a negligible proportion of public or private non-timberlands, in Humboldt and Del Norte Counties, California (Figure 1). The study area was limited to the extent of our high-resolution LiDAR data. Timberland ownership within the 238,355-hectare (588,987-acre) study area was composed of Green Diamond Resource Company (63.8%), other Private (25.0%), State or Federal including Redwood National and State Park, U. S. Forest Service and Bureau of Land Management (8.6%), and Tribal including Yurok and Hoopa lands (1.8%). The remainder (< 1%) was public or private non-timberlands. Hereafter, we will use the term ‘private timberlands’ for both Green Diamond and non-Green Diamond timberlands within the study area. The portion of our study area included in Green Diamond’s HCP application is 145,100 hectares (358,549 acres). Hereafter, we refer to this portion of the study area as the ’Plan Area’.

**Figure 1.**
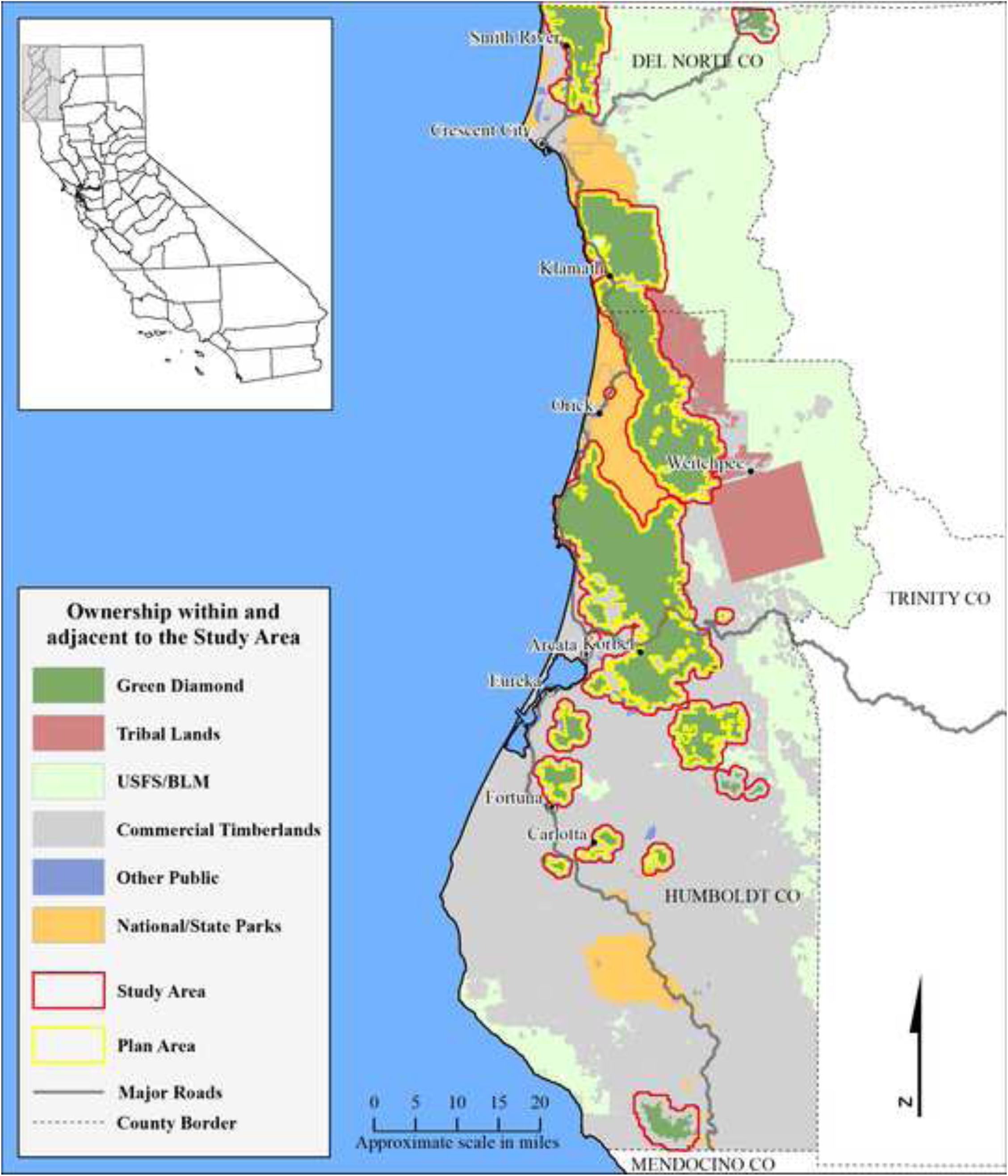
Ownership within the Study Area. The study area, composed of Green Diamond timberlands in Humboldt and Del Norte Counties, and a surrounding 0.5-mile buffer that contained State, Federal, Tribal, other Private Commercial timberlands, and Private non-timberlands. The Plan Area comprised a subset of Green Diamond Timberlands.

The entire study area was within 50 kilometers (18.6 miles) of the Pacific Ocean, and elevation ranged from five to 1550 meters (5,085 ft). The present-day climate on the North Coast of California is characterized as temperate Mediterranean. Along the Pacific Ocean, winters are generally mild and rainy, and summers mild, cool, and dry. Further inland, weather varies from the oceanic climate and temperatures increase throughout the year into a hot summer. Variation in air temperature is generally low along the coast as maritime influence dominates, with decreasing affect inland. Yearly minimum air temperature varies spatially across the study area from -1.6 °Celsius (29.5 °Fahrenheit) to 5.3 ° Celsius (41.6 ° Fahrenheit) with cooler minimum temperatures concentrated inland and in the north. Yearly maximum air temperature varies similarly from 17.5 ° Celsius (63.6 ° Fahrenheit) to 35 ° Celsius (95.0 ° Fahrenheit) with warmer temperatures concentrated inland and in the broad river valleys nearer the coastline. Annual precipitation occurs mostly as rain from November through April averaging 198 centimeters (78 inches) and varies from 106 centimeters (42 inches) near the coast to 320 centimeters (155 inches) in the coastal mountains (PRISM Climate Group, 30-Year Normal data, 1991-2020). Substantial and persistent snow occurs at elevations above 1000 meters (3,280 feet).

In our study area, coast redwood (*Sequoia sempervirens*) forest dominated in the coastal areas and at lower elevations. Douglas-fir (*Pseudotsuga menziesii*) replaced redwood as elevation and distance from the coast increased. Hardwoods exist as pure stands or common forest components. Red alder (*Alnus rubra*) and maple (*Acer macrophyllum*) dominated in coastal mesic sites, while tanoak, madrone (*Arbutus menziesii*) and giant chinquapin (*Chrysolepis chrysophylla*) occurred at higher, more xeric sites.

Private timberlands in the study area have been subjected to timber harvest for more than a century. Forests on private timberlands consist primarily of second and third-growth trees ranging in age from recently harvested to approximately 120 years. Private timberlands contain small and scattered patches of unharvested or partially harvested old growth forest (< 0.2% of private timberlands). Larger intact old growth forests occur on state and federal lands. State and federal lands in the study area consist of some of the largest old growth coastal redwood forests remaining within the California range of the marbled murrelet (Lorenz et al., 2021).

## Methods

We estimated the two-sample HSM (Manly et al., 2002; Warton and Aarts, 2013) for murrelet nesting habitat using Maxent software (Phillips et al., 2004, version 3.4.1). Our two-sample HSM compared environmental conditions (i.e., habitat characteristics) at a sample of ‘used’ locations to those at a sample of ‘available’ locations. We estimated our HSM using the two-sample used-available methodology (Manly et al., 2002), also referred to as presence-background (Phillips et al., 2004; Guillera-Arroita et al., 2015), because we had extensive GIS coverage of known occupied patches but no widespread objective surveys of putative nest trees to establish use and non-use. We selected covariates to include in the HSM using an objective and repeatable four-step procedure. We validated the model by inspecting the magnitude and variation of performance measures over bootstrap replicates drawn from the raw data. We derived habitat classes by objectively identifying changes (“kinks”) in the predicted-expected ratios (Hirzel et al., 2006) over an increasing range of suitability. We estimated concave polygons surrounding the patch and individual trees that influence nesting potential of an area and used these polygons to delineate practical and identifiable stand boundaries and associated habitat quality predictions. These practical and identifiable patch boundaries included trees in the core area of nesting stands but also included peripheral large trees that contributed to nesting suitability.

### Used Habitat Polygons

We constructed polygons of ‘use’ as forest areas with indirect evidence of nesting and labeled them “occupied habitat polygons”. We constructed ‘used’ polygons, rather than used trees or points, due to the paucity of data on known nest locations within our study area. Prior studies faced a similar paucity of nest locations and, like us, defined “occupied behaviors” in a stand as an indication of potential nesting in the stand (Evans-Mack, 2003; Plissner et al., 2015; Washington Department of Natural Resources, 2019). We included as ‘used’ polygons all confirmed occupied patches from Federal and State parks, Bureau of Land Management reserves, private unharvested old growth forests, and private partially harvested old growth within our study area (Miller and Ralph, 1995; Chinnici, 2021; PALCO, 1999). We also included private young (< 70 years old) forests with residual old growth trees containing “occupied behaviors”.

On private timberlands, we established murrelet occupied behaviors in forest stands and contiguous habitat by three methods. We established occupancy using audio-visual surveys conducted according to inland survey protocols for marbled murrelets that account for imperfect detection (Ralph et al., 1994; Evans et al., 2000; Evens Mack et al., 2003), using radar surveys (Bigger et al., 2006), and using direct evidence of murrelet nesting reported by other studies (Hebert and Golightly, 2006; Golightly et al., 2009). The audio-visual inland survey protocol on private timberlands dictated multiple site visits (≥ 5) over a minimum of two years. During these surveys, we noted occupied behaviors such as stationary vocalizations from a tree, landing in a tree, flights below and through the forest canopy, and circling flights below or above the forest canopy. If we observed murrelet occupied behavior, we assigned ‘used’ status to the area. If we failed to observe occupied behaviors, the forested areas were cleared for harvest under California timber harvest plan permits (California Department of Forestry and Fire Protection, 2024). Hence, the audio-visual surveys provided direct evidence of non-use (absence) when we failed to observe occupied behavior. Unfortunately, many of these non-use areas were subject to timber harvest prior to acquisition of the LiDAR data and we could not derive accurate LiDAR covariates in these areas. We included these cleared areas among the ‘available’ habitat.

When we observed occupied behavior, we established the boundaries of the patch that contained the occupied behavior based on notable differences in forest age (or structure) such as between unharvested or lightly harvested old growth habitat (> 200 years old) and adjacent young managed forests generally less than 70 years old. We also considered the concept of contiguous habitat from surveyed areas (Evens Mack et al., 2003). On Green Diamond timberland, we included 24 patches that contained historic indirect evidence of nesting, and three patches with recent indirect evidence of nesting. We established the boundaries of the three recent patches using spatial kernel density estimation (95% KDE) of audio/visual detections.

### Balanced Sampling – ‘Used’ and ‘Available’ Points

After establishing ‘used’ polygons, we drew a single balanced acceptance sample (BAS) (Robertson et al., 2013, 2013; Brown et al., 2015) containing 117,804 locations across the entire 238,310-hectare (588,838-acre) study area. Our BAS had an average density of one sample point per 2.02 hectares (5-acres). We extracted all BAS points falling inside ‘used’ polygons, but then excluded those within 20 meters of their patch boundary. We removed points inside but near the patch boundary to reduce edge effects (bias) in LiDAR-derived moving window covariates. We chose the inner exclusion zone distance of 20 meters, which is less than the 35.89-meter radius of our smallest 1-acre moving window, to maximize the number of ‘used’ locations while balancing our concern over edge effect bias. We identified a total of 1,080 BAS points falling inside the ‘used’ polygons after excluding those within 20 meters of patch boundaries (Figure 2). We considered these 1,080 points as ‘used’, while we considered the totality of BAS points (117,804 points) as ‘available’.

**Figure 2.**
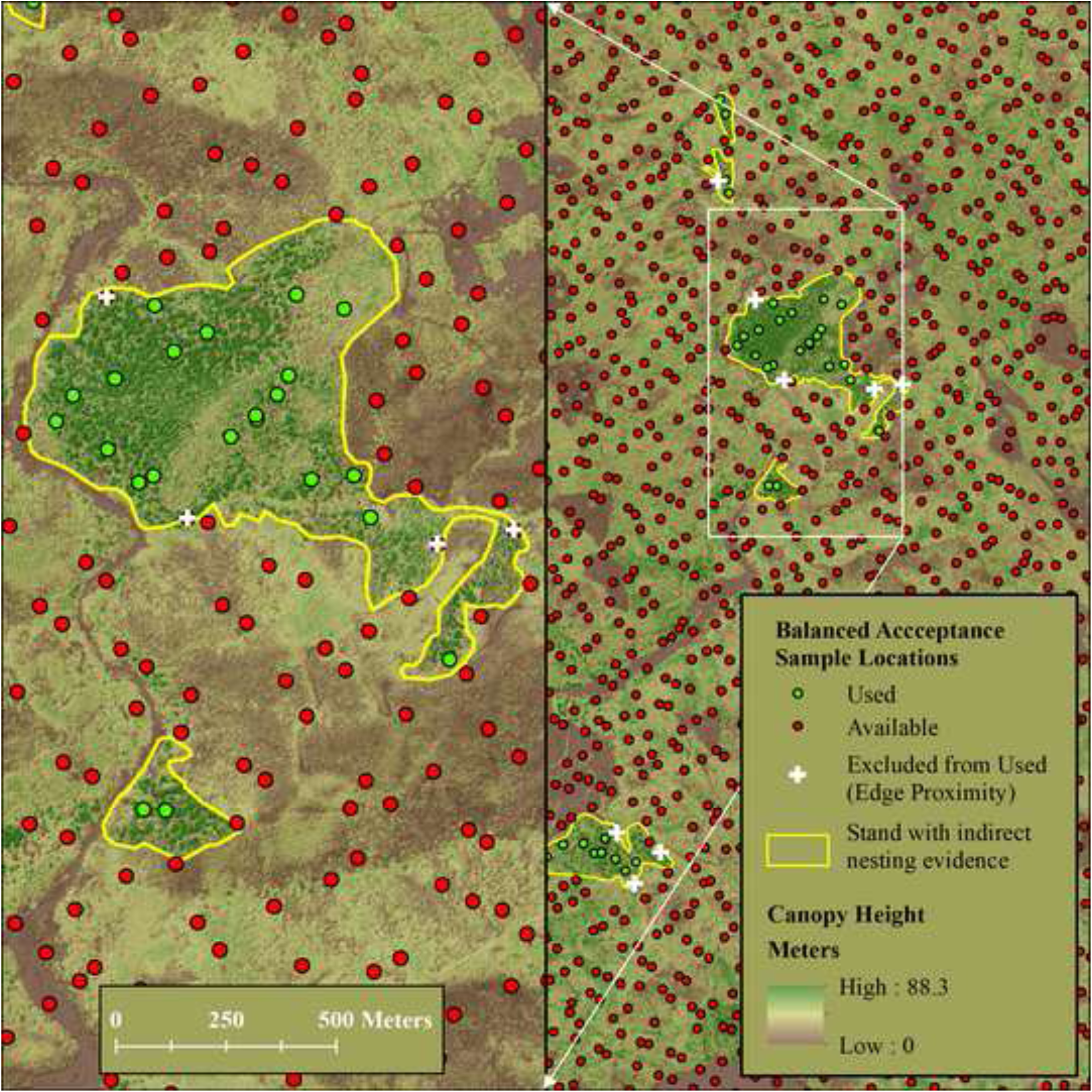
Balanced acceptance sampling (BAS). Illustration of Balanced Acceptance Sample (BAS) within the study area. We sampled ‘used’ points (green dots) and ‘available’ locations (both green and red dots) at an average density of one point per 5 acres. We excluded from the sample of ‘used’ points any location within 20 meters inside the boundary of a patch with indirect evidence of nesting to reduce moving window edge effects (white crosses). We classified all BAS points (inside, outside, and near boundary) as ‘available’. Dark brown areas with low canopy height are prior clear cuts of varying age.

### LiDAR Data

LiDAR data was collected from December 2017 through February 2018 using a Cessna Caravan aircraft carrying a Riegl VQ-1560i LiDAR sensor. Wave form data was collected and discretized in real-time on board the aircraft yielding up to 14 returns per pulse. Survey altitude above ground (AGL), speed, swath overlap, and central wavelength of the laser were 1,306 meters, 120 knots, 55%, and 1064 nanometers, respectively. Specifications called for a minimum density of 30 pulses/m^2^. Our realized pulse density averaged greater than 40 pulse/m^2^, with a maximum of 560 returns per m^2^. Ninety-five percent of returns had horizontal and vertical accuracy of ≤ 20 centimeters. Raw LiDAR data consisted of the rectified and classified 3-dimensional location of every return collected by the sensor.

We processed the raw LiDAR data in LASTools (2019) which among other things allowed us to discretize the (*x, y, z*) point cloud into ‘voxels’ (a 3-dimensional pixel extending from ground to AGL), measure the distribution of points inside voxels, and develop a 0.25-meter resolution Digital Surface Model (DSM) for the canopy surface. We derived Tree Approximate Objects (TAOs) by applying a local maxima process to the canopy DSM that ultimately extracted the (*x, y*) location and height (*z*) of the TAO (Popescu et al., 2004; Tiede et al., 2005; Chen et al., 2006; Jeronimo et al.; 2018).

### Covariates

We compiled eighty HSM covariates on a 5-meter grid covering the entire study area. We grouped these eighty covariates into six general themes (Hirzel and Le Lay, 2008) based on their hypothesized importance as predictors of murrelet nesting habitat (Table S1). The ‘Biographic/Geographic’ group included 14 covariates such as elevation, distance to coast, aspect, and solar radiation. The ‘Climate” group included 6 covariates such as annual precipitation, air temperature, growing degree days, and whether a location was in the zone of coastal influence. The ‘LiDAR Canopy Metrics’ group included 10 canopy metrics including the number of returns in each 5-meter by 5-meter voxel footprint, average return height above ground (a measure of central canopy height), and variation in returns within a given voxel. The ‘LiDAR Percentile’ group included 19 variables measuring the height inside each voxel from which a certain percentage of the returns were received. We derived 16 TAO based statistics in the ‘TAO Point Statistics’ group such as the minimum, mean, maximum, sum and standard deviation of TAOs ≥ 50 meters in height within 1- and 5-acre circular plots surrounding the (*x, y*) location of a voxel (North et al., 2017). The ‘TAO Point Density’ group included 15 covariates such as number of TAOs ≥ 50 meters within a 1- and 5-acre circular plot centered on the voxel’s (*x, y*) location. Only covariates in the ‘TAO Point Statistics’ and ‘TAO Point Density’ groups were averaged over spatial areas and subject to bias near patch edges, which we mitigated by eliminating used points within 20 meters of patch boundaries. Spatial reference for our 5-meter resolution covariate grids was NAD83, UTM Zone 10.

### Model Selection

We investigated pairwise correlations among covariates by computing Pearson’s R^2^, Cramer’s V, and box plots. From these, we identified pairs and groups of correlated variables that could cause model instability if included in the same model. We identified correlated covariates despite the known robustness of two-sample HSMs to covariate collinearity (Elith et al., 2011; Feng et al., 2019). We considered these sets of correlated covariates during all phases of model selection, except the first.

We performed HSM model selection in 4 phases. Phase 1 screened covariates in each of our six covariate themes by fitting all theme covariates to the ‘used’ and ‘available’ points at once in Maxent. Strongly correlated covariates in the same group were allowed in the Phase 1 model because Phase 1 simply screened variables based on predictive strength. We assessed the predictive strength of individual covariates using a combination of measures reported by Maxent. We evaluated marginal plots, percent contribution, permutation importance, regularized training gain, and testing gain to assess predictive strength (Phillips, 2017). High strength covariates exhibited a biologically reasonable marginal response and high relative influence (typically values ≥ 20), as measured by percent contribution and permutation importance. Maxent’s jackknife test measured the goodness of fit of the model. We advanced covariates with jackknife values of approximately 2.5 or better during both regularized training and testing gain. A jackknife value of 2.5 translates to a likelihood improvement of 12.2 (= exp (2.5)) times relative to the null model (Young et al., 2011).

During phase 2, we combined the top Phase 1 covariates across groups into a single model but prohibited correlated covariates from appearing in the same model. We defined covariates to be correlated when their pairwise Pearson and Cramer V statistics exceeded 0.65. Due to these correlation prohibitions, we ultimately obtained two HSM models each containing nine pairwise uncorrelated covariates. At the end of Phase 2, we advanced the covariates from each model that exhibited high predictive strength based on percent contribution, permutation importance, regularized training gain, and testing gain.

During Phase 3, we divided the five unique covariates that advanced from Phase 2 into four groups of three uncorrelated covariate groups (Pearson’s R^2^ less than 0.65). We fitted each of the four models and assessed AIC (Akaike, 1974) over 10 bootstrap resamples of the data. We inspected boxplots of AIC over bootstrap iterations and advanced to Phase 4 the model with the lowest median AIC.

During phase 4, we optimized Maxent’s regularization multiplier using the best Phase 3 model. Prior phases had Maxent’s regularization multiplier set to 2. Phase 4 was designed to investigate the predictive strength of multipliers around 2. We tried nine regularization multipliers from 1.0 to 5.0 in steps of 0.5 and chose the value with lowest median AIC over 10 bootstrap resamples.

During all model selection phases, we set Maxent to perform 10 bootstrap replicates each with 5000 fitting iterations. We used scaled logistic output and model form combinations of Linear, Quadratic, Product, and Hinge shapes. During each bootstrap replicate, Maxent fitted the HSM model to a random subset of 80 percent of the selected ‘used’ locations and computed model fit statistics on the remaining 20 percent.

### Habitat Classification

To classify habitat across the study area, we estimated break points in a smoothed continuous P/E ratio curve (Boyce et al., 2002; Hirzel et al., 2006). We constructed the P/E curve from our final HSM predictions using overlapping bins of predicted HSI values with widths equal to 0.02 centered on values from 0.02 to the maximum prediction, in steps of 0.002. Segments of the P/E curve with different slopes indicate changes in the accumulation of ‘used’ points relative to ‘available’ and represented a change in selection habits relative to the other segments (Figure 3). To identify specific breakpoints, we fitted a piece-wise linear curve via ordinary least-squares to the smoothed P/E curve and searched for the best fitting habitat breakpoints via direct search across all possible breakpoint combinations. We computed breakpoints on a smoothed version of the raw P/E ratios that we smoothed using function ‘`supsmu`’ in R (Friedman, 1984) to aid visual interpretation and speed the search for breakpoints; but the identified breakpoints varied little when we segmented the un-smoothed P/E curve.

**Figure 3.**
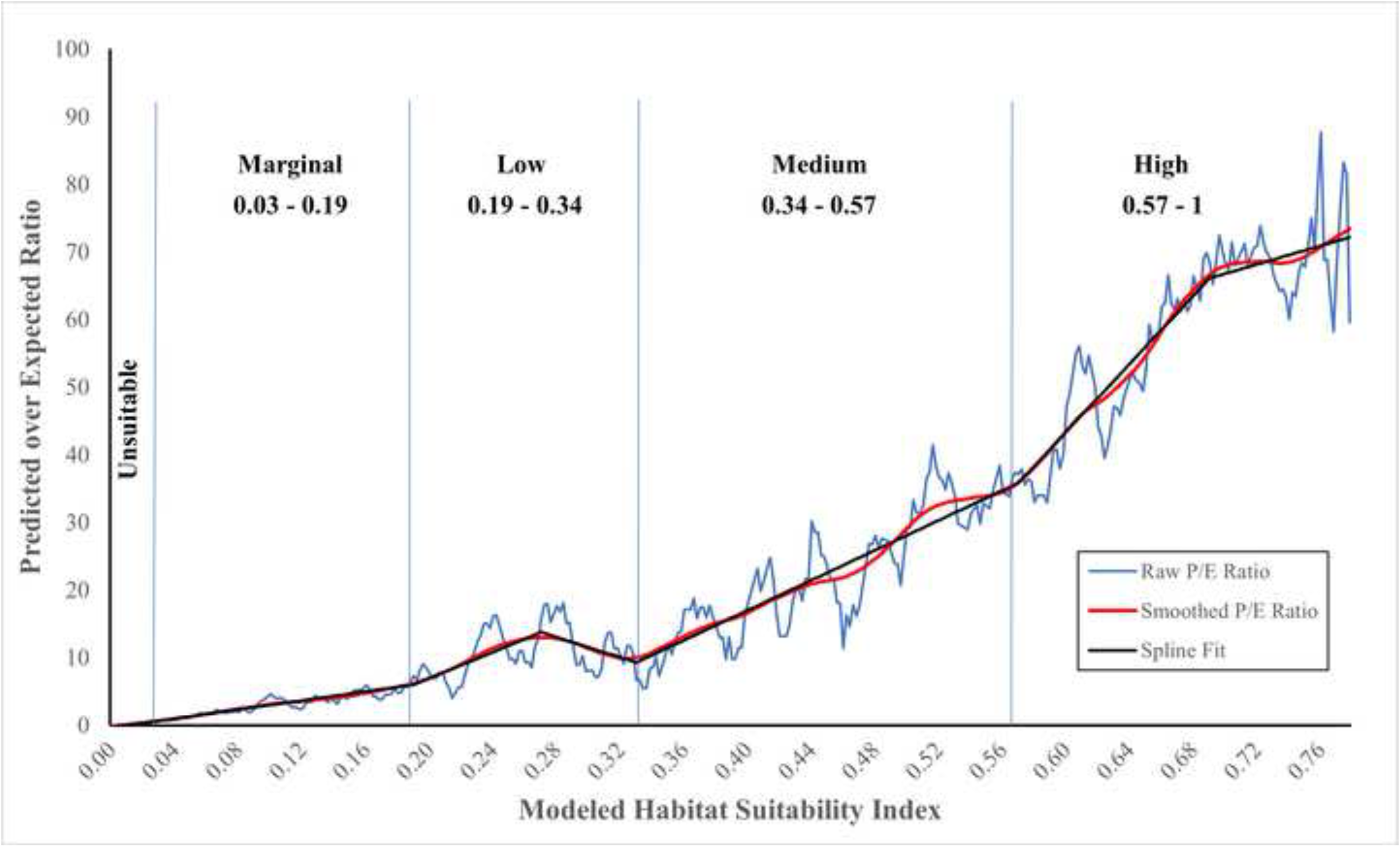
Predicted-to-Expected (P/E) ratio curve. P/E curve from the final HSM model showing the continuous Boyce (raw P/E ratio) curve, smoothed curve (Friedman 1984; function supsmu in R) and fitted lines from a spline model with kinks at habitat classification breaks.

We forced the first habitat breakpoint at the maximum HSI value with P/E < 1.0 because those areas contained fewer ‘used’ points than expected by chance (Hirzel et al., 2006). We regarded locations with P/E < 1.0 to be “unsuitable”. For P/E values > 1.0, we combined areas with predicted values between each of the best-fitting breakpoints and attached a descriptive label to the class. Ultimately, we identified four habitat classes for areas with P/E values > 1.0 that we labeled “marginal”, “low”, “medium”, and “high” (see Results). We used these categories to delineate murrelet habitat patches.

### Patch Delineation

We identified marbled murrelet habitat patches by first mapping habitat classes onto 5-meter raster cells (i.e., pixels) covering the entire study area. We then buffered patches of “low” or better habitat outward by 80.25 meters (263 feet) to encompass areas adjacent to the patch that could have contributed to habitat quality inside the “low” and better patch (Figure 4). Our buffer distance (i.e., 80.25 meters) was the radius of the 2-hectare (5-acre) circular moving windows that were a part of the best model’s TAO-based covariates and hence TAOs within this distance contributed to predictions inside the patch. Within this buffered region, we identified all TAOs ≥ 50 meters. We then placed a concave hull (Park and Oh, 2012) around all TAOs ≥ 50 meters and smoothed its perimeter to construct boundaries around patches of habitat with better-than-random selection characteristics. We computed concave hulls using the `concaveman` R package (Gombin et al., 2020) with concavity parameter set to 0.8 and default length threshold set to 0.0. We smoothed the perimeter of our concave hulls by computing a 60-meter outward buffer followed by a 55-meter inward buffer. This outward and inward buffering process removed many internal intrusions (bays between peninsulas) when they occurred and thereby made the perimeter both more conservative (larger), realistic, and easier to follow in the field.

**Figure 4.**
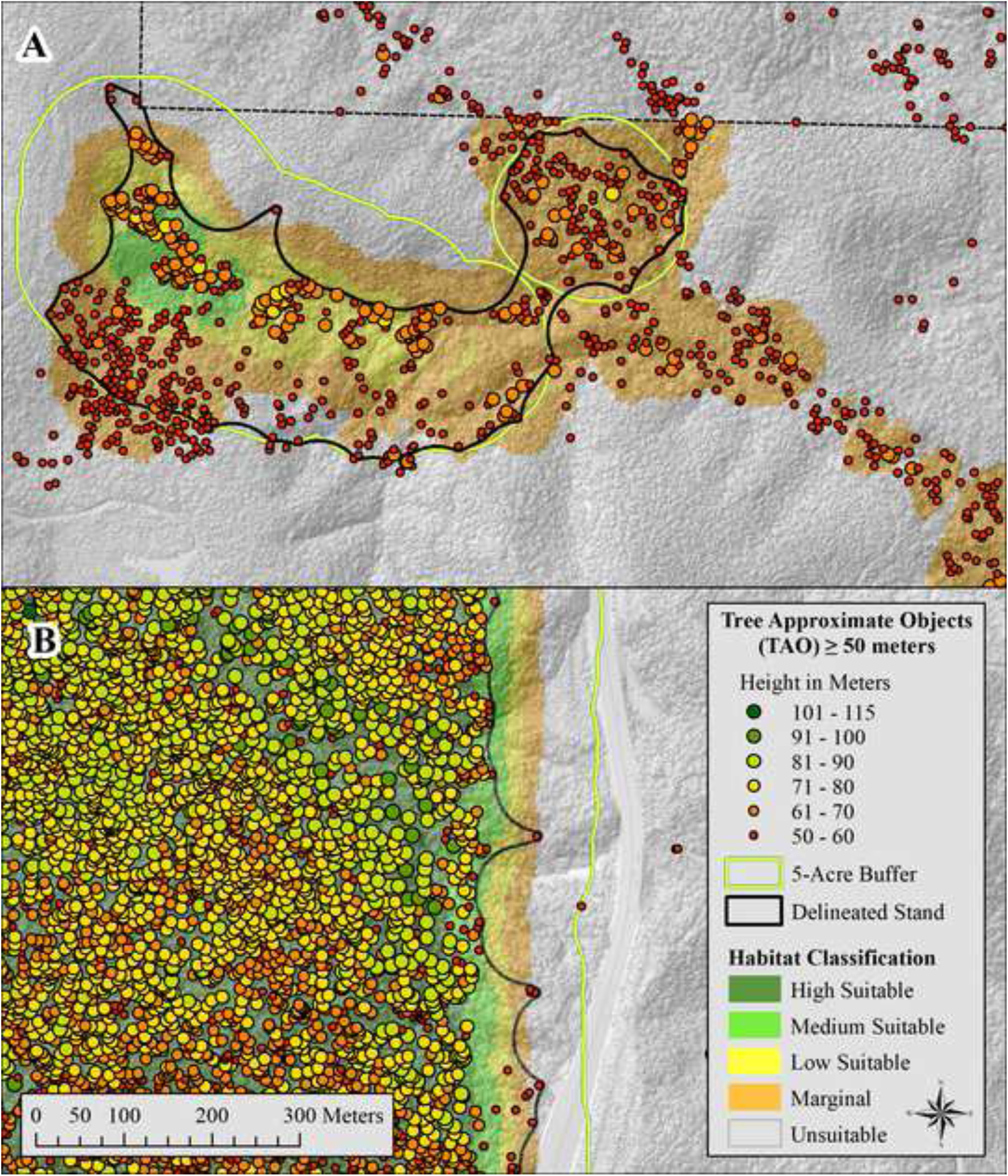
Habitat delineation and TAOs. Map panels showing examples of delineated marbled murrelet habitat patches. Patch boundaries (black) were derived by constructing a five-acre (2-hectare, 80.25 meter) buffer around Low or better suitability pixels (HSI values ≥ 0.19, Figure 3), selecting all tree approximate objects (TAO) ≥ 50 meters in height inside the buffer, and computing a concave hull around these trees. TAOs ≥ 50 meters within 80.25 meters of a pixel contributed to predictions of Low and better habitat areas. Panel (A) illustrates a higher quality patch delineated on Green Diamond property within the Plan Area and panel (B) illustrates an area of old-growth forest in Redwood National and State Park between the Plan Area boundary and the ½ mile outer buffer delineating the boundary of the Study Area.

### Model Evaluation

We used the receiver operating characteristic curve (ROC) and area under curve (AUC) output from Maxent to evaluate overall model prediction ability. AUC ranges from 0 to 1, with a value of 0.5 indicating predictions no different than random chance and with AUC = 1 indicating perfect discrimination. In addition to AUC, we computed the sensitivity of our model to correctly predict ‘used’ locations given a specific threshold above which locations were considered “habitat” and below which locations were considered “not habitat”. Using that same threshold, we computed the Positive Predictive Value (Parikh et al., 2008) and Cohen’s Kappa (Cohen, 1960; Smeeton, 1985) to assess the predictive value of our HSM. We evaluated marginal plots of relative probability of selection (i.e., HSI value) by each covariate in the final model to gain insight into model predictions and to assess the biological reasonableness of predictions in each dimension.

## Results

### HSM Model Selection

The six Maxent runs completed during Phase 1 (covariate screening) yielded eleven covariates (Table S2). We advanced three covariates from the TAO Point Density and TAO Point Statistics themed groups, two from the Biogeographic/Geographic group, and one each from the TAO Percentiles, Canopy metrics, and Climate groups. At the end of Phase 1, we discounted two covariates (Coast_Dist and PPT) with high influence because they exhibited implausible bimodal marginal plots at the extremes of their ranges. We ultimately discounted another variable (ZCI) that was initially elevated to Phase 2 but exhibited very low Permutation Importance and regularized training/testing gain. The categorical nature of the ZCI variable, essentially defining the coastally influenced fog zone, provided little spatial discrimination in our study area.

After separating correlated covariates, we fitted two Maxent models during Phase 2 of model selection (Table S2). Both models contained nine uncorrelated variables. We identified top variables in both models, removed duplicates among the top models, and advanced five covariates from Phase 2 to 3. All five advancing covariates came from the TAO-based density and height themed groups. Covariates in the percentile and canopy metric groups, as well as those in the biogeographic, and climate groups displayed little predictive value compared to the TAO-based density and height covariates. In this stage, Simpson’s Diversity Index of TAO height classes (SDI_3cl_5ac) appeared to perform well exhibiting a high Percent Contribution but yielded a very low Permutation Importance. During ancillary Phase 2 investigations, we discovered computational inconsistencies in the Simpsons’ Diversity results and dropped it.

During Phase 3, we fitted four models that represented all possible models of uncorrelated covariates (Table S2). Among these four, the model containing density of TAOs ≥ 60 meters, total height of TAOs ≥ 50 meters, and the standard deviation of TAO heights ≥ 50 meters produced the lowest median AIC (ΔAIC to next best model = 39.3). This model also produced the least variability in AIC among the four models tested (Figure 5), suggesting stability and consistency among random replicates. Among covariates in this model, tree density (TAOs ≥ 60 meters) and total height (of TAOs ≥ 50 meters) contributed more to predictions than the standard deviation of tree height (of TAOs ≥ 50 meters).

**Figure 5.**
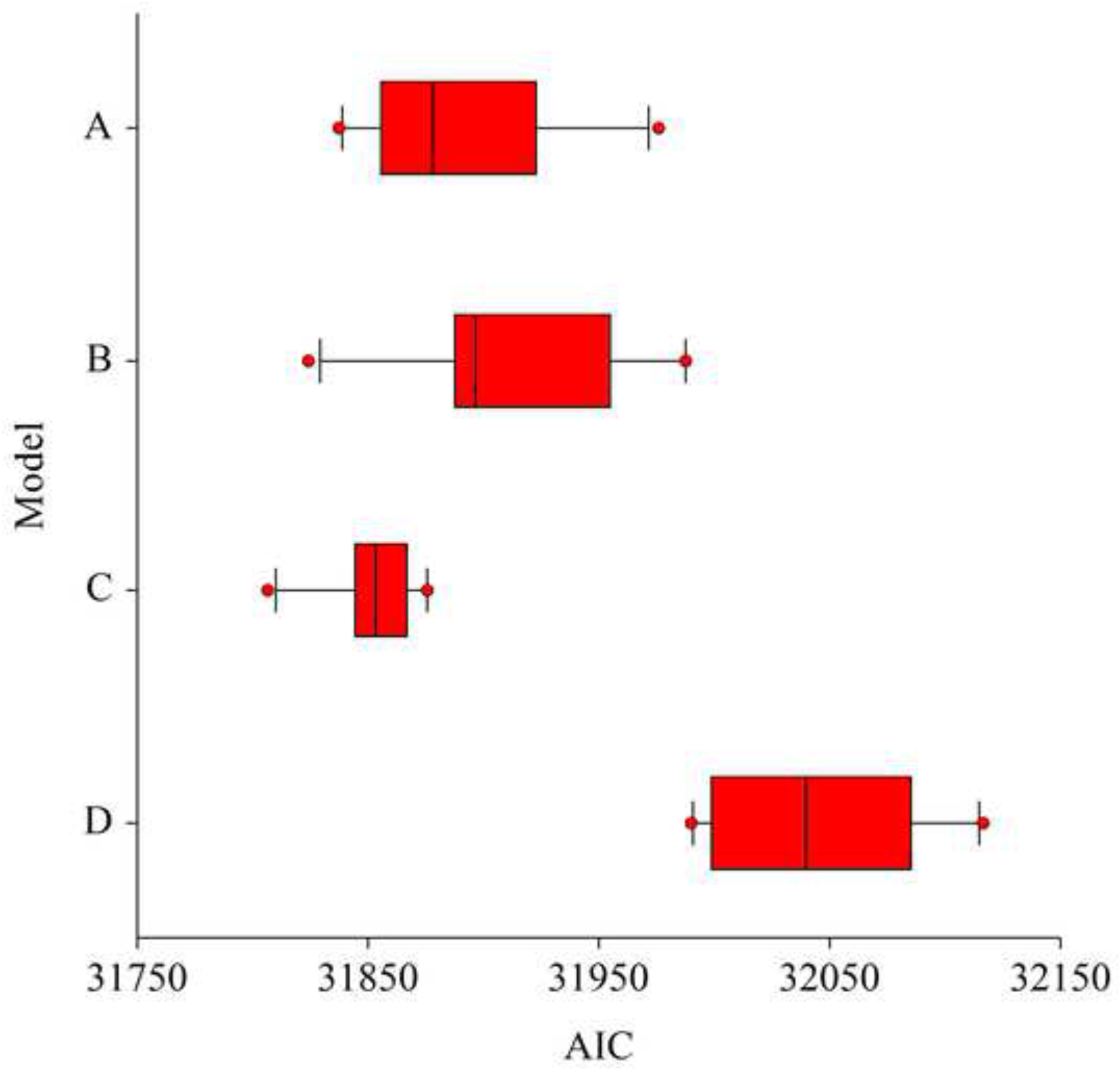
AIC plot - Final model selection. Box and whisker plots of Akaike Information Criteria (AIC) values for the four models with three-variable combinations. The best three-variable model (C) consisted of the point density of tree approximate objects (TAO) ≥ 60m within a 5ac buffer, the sum of heights of TAO ≥ 50m within a 5ac buffer, and the standard deviation of TAO ≥ 50m within a 5ac buffer. Median and dispersion for model C are the lowest suggesting both parsimony and consistency between replicates. Box boundaries are at the 25th and 75th percentiles, the whiskers extend from the 10th to the 25th and the 75th to the 90th percentiles, and the outliers (red dots) are represented beyond the whiskers.

We ran Maxent nine times during Phase 4 to identify the best regularization multiplier (RM) between 1.0 and 5.0. The Maxent run with RM equal to 2.0 produced the lowest AIC among those tested and the least amount of variation over bootstrap replicates (Table 1, Figure 6). We set RM to 2.0 during covariate selection (Phases 1 through 3) and we were not surprised that RM = 2.0 produced the lowest AIC because we found in prior Maxent modeling efforts that an RM = 2.0 is a good compromise between model complexity (lower RM values) and simplicity (higher RM values). We acknowledge that use of a different RM during Phases 1 through 3 could conceivably have resulted in a different best-fitting model. Regardless, the minimum AIC with RM = 2.0 supports the idea that no other combination of RM and covariates in our final model performed better, and subsequent model evaluation indicated that our final model adequately fitted the ‘used’–‘available’ data.

**Figure 6.**
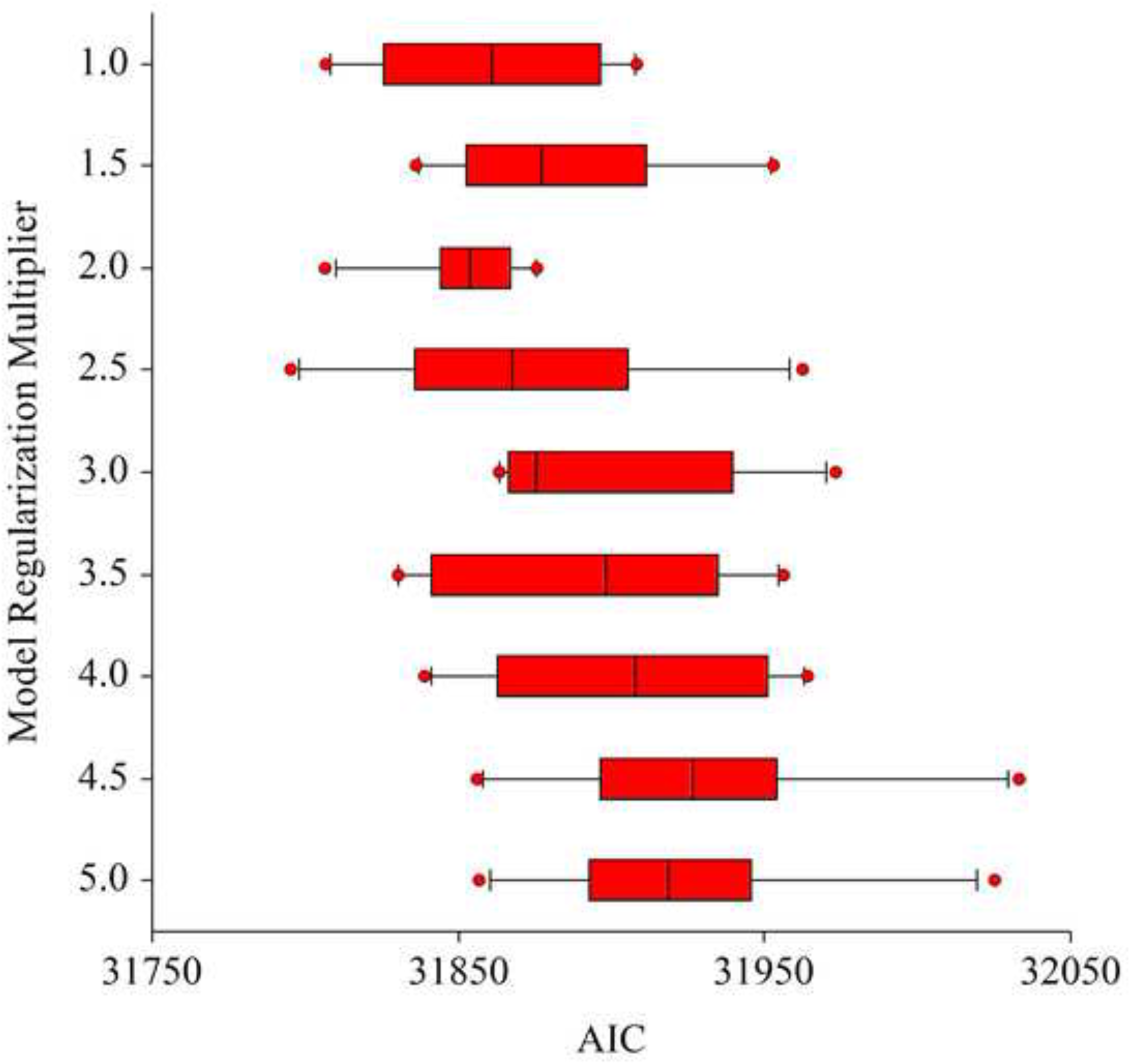
AIC plot - Regularization Multiplier assessment. Box and whisker plots of Akaike Information Criteria (AIC) values for the best three-variable model (C) with regularization multiplier varied from 1.0 to 5.0 in 0.5 increments. Median and dispersion for the model with RM = 2.0 are the lowest suggesting both parsimony and consistency between replicates. Box boundaries are at the 25th and 75th percentiles, the whiskers extend from the 10th to the 25th and the 75th to the 90th percentiles, and the outliers (red dots) are represented beyond the whiskers.

**Table 1.** AIC based Regularization Multiplier assessment. The results of nine iterative replicated model runs of the best candidate (3var_C) resulting from varying the Regularization Multiplier (RM) in Maxent from 1 to 5 by steps of 0.5 and listed by Akaike Information Criteria (AIC) values. The RM value of 2.0 exhibited both the lowest AIC value and least AIC dispersion between replicates followed by a non-competitive RM of 1.0 (ΔAIC of 6.8)

| Regularization Multiplier | Average Number of Model Parameters | Average of log Likelihood | Average AIC | Delta AIC |
| --- | --- | --- | --- | --- |
| 2.0 | 36.7 | -15889.3 | 31852.0 |  |
| 1.0 | 42.7 | -15886.7 | 31858.7 | 6.8 |
| 2.5 | 34.7 | -15900.5 | 31870.5 | 18.5 |
| 1.5 | 38.7 | -15902.9 | 31883.3 | 31.3 |
| 3.5 | 30.0 | -15916.7 | 31893.4 | 41.4 |
| 3.0 | 33.5 | -15913.8 | 31894.7 | 42.7 |
| 4.0 | 34.3 | -15918.1 | 31904.7 | 52.7 |
| 5.0 | 30.2 | -15932.0 | 31924.5 | 72.5 |
| 4.5 | 33.6 | -15931.6 | 31930.4 | 78.5 |

Marginal plots for the three TAO-based density and height covariates in our final model (Figure 7) revealed positive correlations between relative probability of selection and all three measures. HSI values associated with TAO density increased linearly from 0 (no trees ≥ 60 meters) to 25 tall trees per hectare (10 trees per acre) in a surrounding 2-hectare (5-acre) circle. HSI values associated with TAO height standard deviation increased linearly from 0 to 6 meters (19.7 feet) of large tree height standard deviation in a 2-hectare (5-acre) circle, respectively. Beyond those levels, relative probability of selection plateaued and did not increase appreciably. HSI values increased curvilinearly and plateaued as total height (sum of heights) of large trees in the surrounding 2 hectares (5 acres) increased from 0 to 15,000 meters.

**Figure 7.**
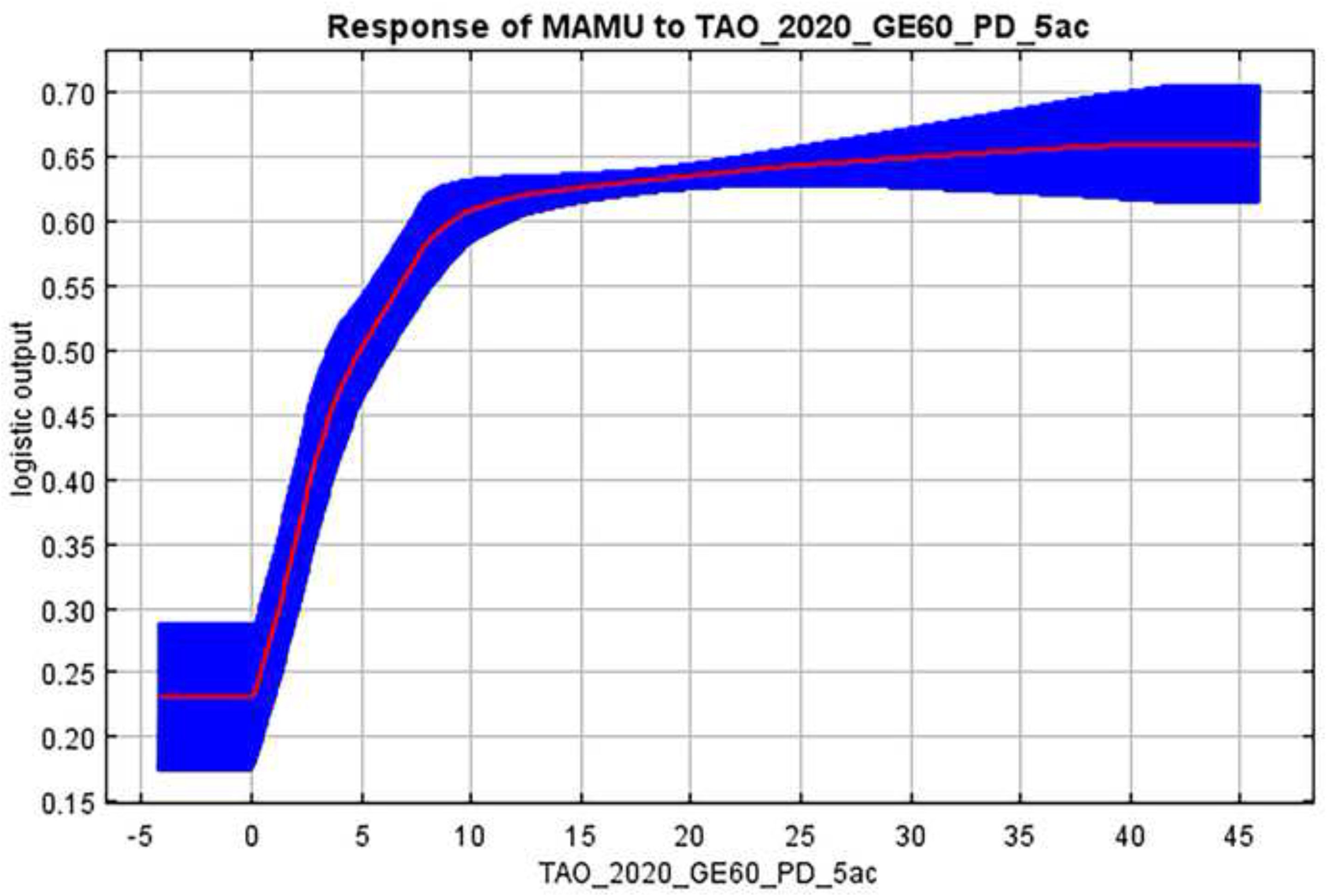
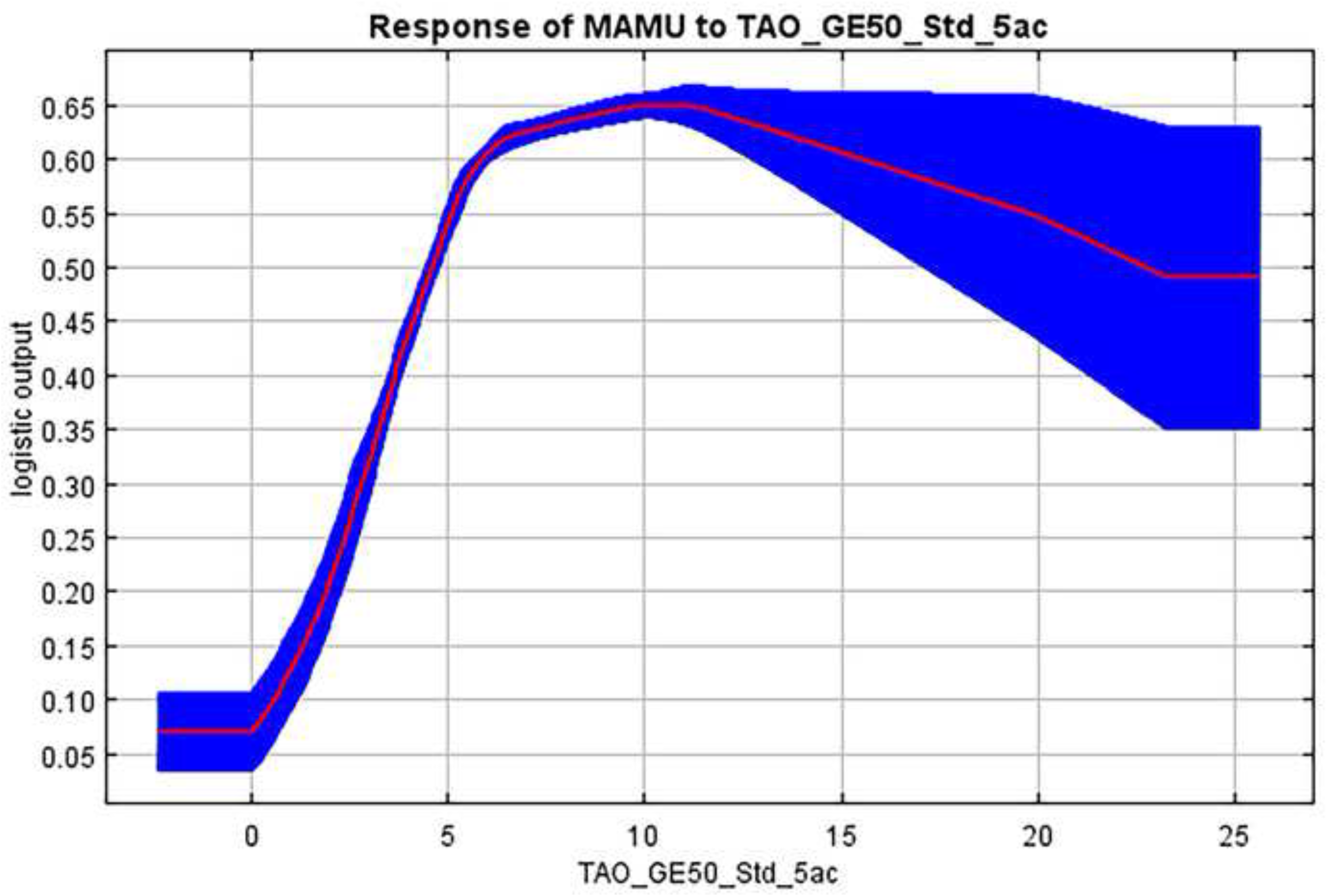
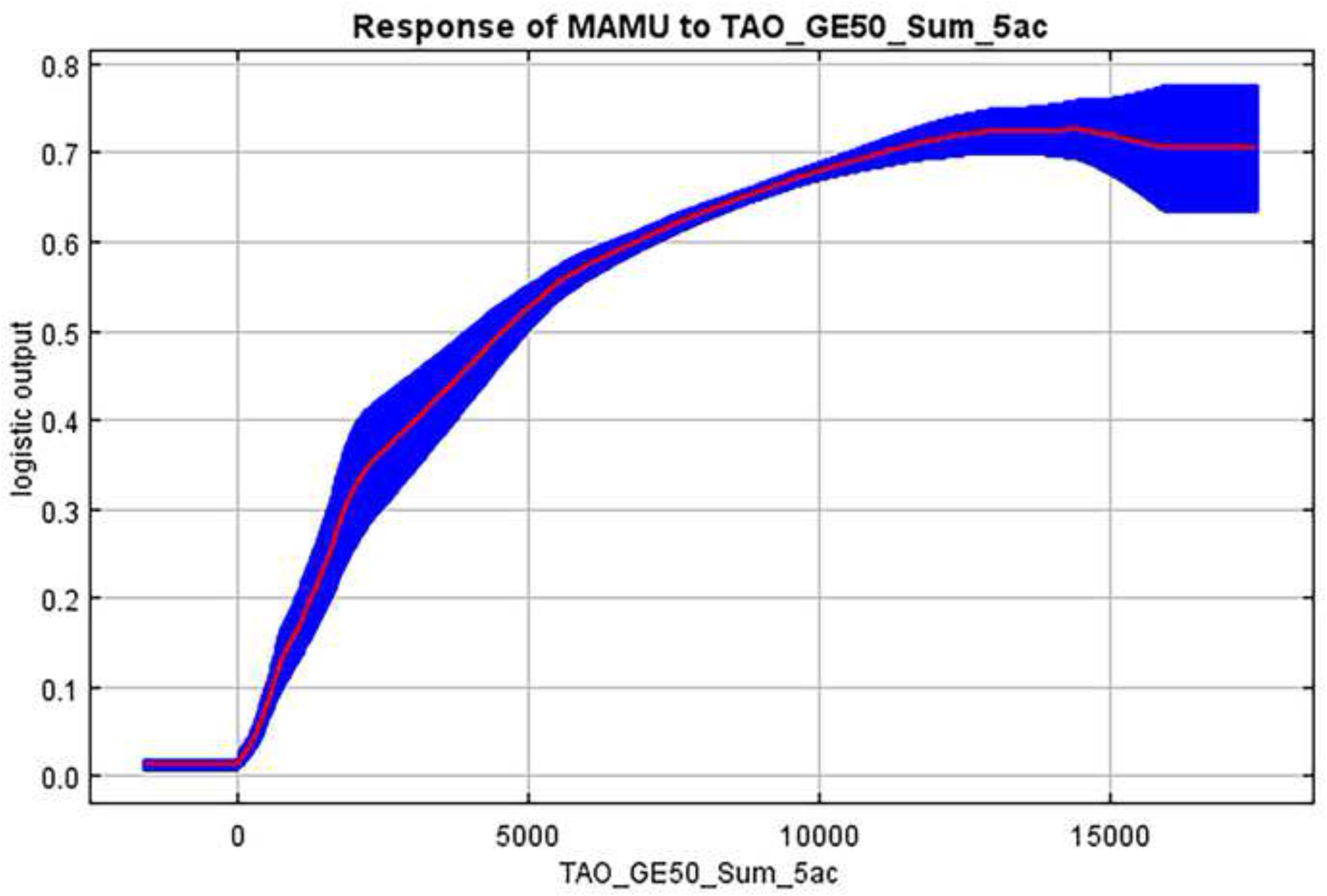
(A, B, and C) Final model covariate response plots. Marginal plots showing response of relative habitat suitability to final model covariates: (A) Density per acre of TAOs ≥ 60 meters tall within a 5-acre circle, (B) Standard deviation in height for TAOs ≥ 50 meters tall within a 5-acre circle, and (C) Sum of heights for TAOs ≥ 50 meters tall within a 5-acre circle. The red line represents the mean value for 10 replicate runs and the blue shade is ± one standard deviation. Units of x-axis values are trees per acre.

### Habitat Classification

The continuous P/E curve for our study showed a generally increasing trend from 0.0 up to the maximum HSI value of 0.78 (Figure 3). Our P/E ratio exceeded 1.0 for HSI values greater than 0.03; hence we considered areas with HSI <= 0.3 as “unsuitable” The best fitting break point model for HSI values above 0.03 included five break points at 0.19, 0.27, 0.34, 0.57, and 0.69. The P/E segment between 0.27 and 0.34 identified P/E ratios with negative slope in HSI value. Decreasing P/E on this segment may be due to removal of ‘used’ locations within 20 meters of ‘used’ patch edges that we implemented to reduce edge effects. We consider the brief and small decrease in P/E on this segment an anomaly in an otherwise monotonically increasing curve and immaterial to management implications; consequently, we removed the 0.27 breakpoint thereby combining the adjacent categories. In addition, we discarded the breakpoint at 0.69 because it fell at the highest end of the suitability curve where the slope of the line flattens and hence imparted little management concern. The final set of breakpoints delineated the five habitat classes which we labeled unsuitable (0 – 0.03), marginal (0.03 – 0.19), low (0.19 – 0.34), medium (0.34 – 0.57), and high (0.57 – 0.78). We note that a large proportion of the “marginal” areas were adjacent to and dependent upon areas of “low” or better habitat. That is, without the supporting TAOs in nearby “low” or better habitat, a large proportion of our “marginal” habitat would have been deemed “unsuitable” (Figure 4).

Under our classification scheme, the study area, and smaller Plan Area were heavily dominated by unsuitable habitat (96.01% and 98.08% respectively) due to the scarcity of TAOs ≥ 50 meters (Table 2). The remainder of the Study and Plan Areas are mostly marginal (2.36% and 1.63%), with less than 0.7% in the low, medium, or high classifications. We found that 20 of the 24 historic known used patches in the Plan Area contained at least one HSI value in the medium or high habitat suitability index range, while 17 of the 24 contained average HSI values in the low or better classes (Figure 8).

**Figure 8.**
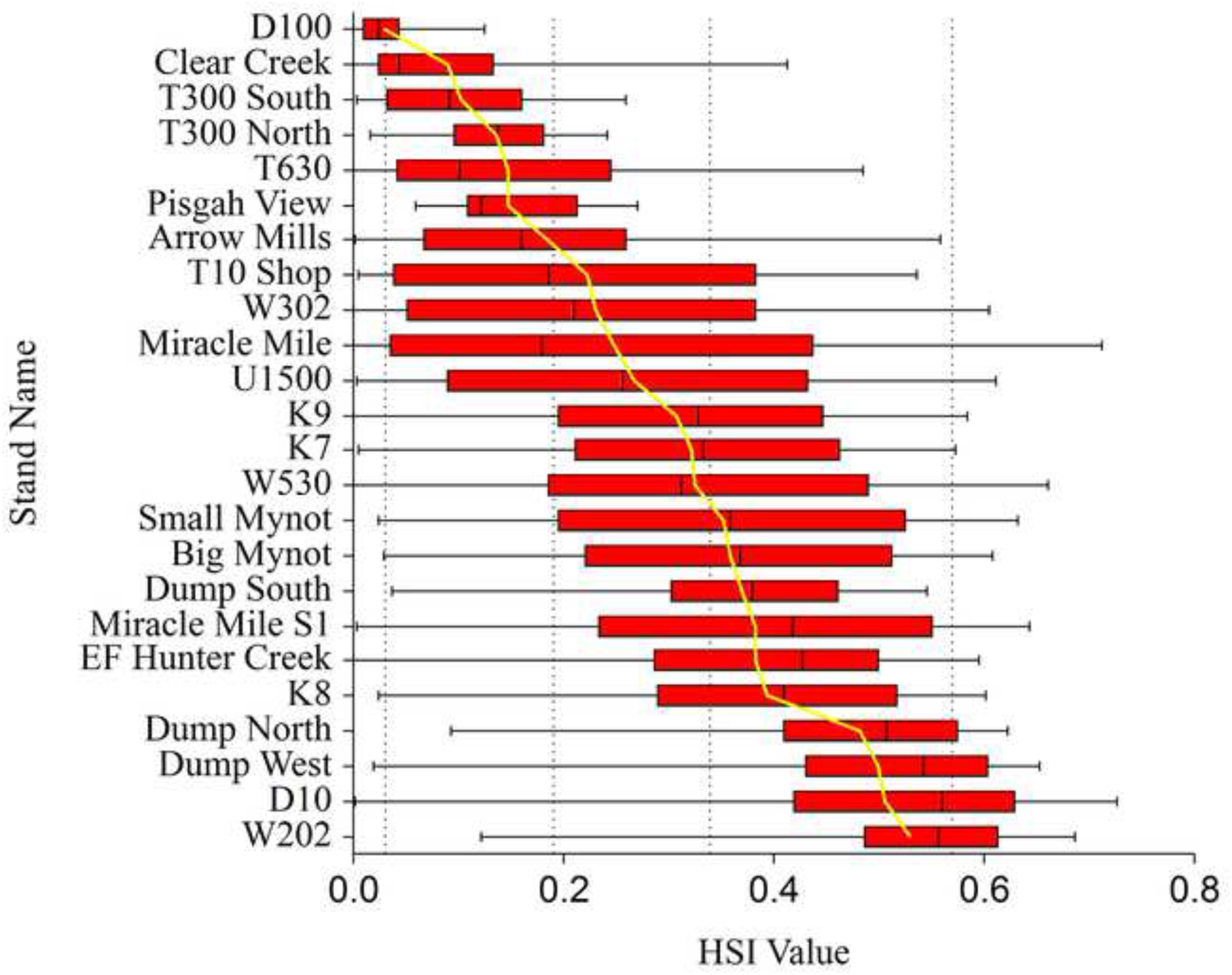
Mean and median HSI values for Historic Murrelet Stands. Box and whisker plots showing mean and median Habitat Suitability Model (HSM) values for 24 historic forest stands containing observed marbled murrelet nesting behavior. Box boundaries are at the 25th and 75th HSM percentiles, the whiskers extend to the extents of the range, and the yellow line represents the mean HSM value. Vertical dashed lines represent the boundaries of the derived habitat classification; unsuitable (0 - 0.03), marginal (0.03 - 0.19), low (0.19 - 0.34), medium (0.34 - 0.57), and high (≥ 0.57).

**Table 2.** Summary of habitat classes by Plan and Study Area. Areal extant and percentage within each marbled murrelet habitat class (HSM raster), along with relative habitat suitability index (HSI) range, on the Plan and Study area.

| Habitat Class | HSI Range | Plan Area |  | Study Area |  |
| --- | --- | --- | --- | --- | --- |
|  |  | Hectares (Acres) | Percent | Hectares (Acres) | Percent |
| Unsuitable | 0.0 – 0.03 | 142,319 (351,678) | 98.08% | 228,847 (565,491) | 96.01% |
| Marginal | 0.03 – 0.19 | 2,370 (5,856) | 1.63% | 5,617 (13,880) | 2.36% |
| Low | 0.19 – 0.34 | 243 (601) | 0.17% | 1,106 (2,732) | 0.46% |
| Medium | 0.34 – 0.57 | 143 (354) | 0.10% | 1,155 (2,855) | 0.48% |
| High | 0.57 – 0.78 | 24 (60) | 0.02% | 1,630 (4,028) | 0.68% |
| Total |  | 145,100 (358,549) | 100.00% | 238,355 (588,987) | 100% |

### Patch Delineation

In the Study Area, we identified 312 patches encompassing 5,421 hectares (13,395 acres; Figure 9). Similarly, within the smaller Plan Area, we identified 171 patches, encompassing 729 hectares (1,802 acres), that received classifications of low suitability or better (Table 3). Within the Plan Area, twenty-four of the 171 patches occurred in or adjacent to known ‘used’ patches. An additional 34 patches in the Plan Area were near boundaries where most of the contributing TAOs were located just outside of the Plan Area but still in the Study Area (e.g., Prairie Creek Redwoods State Park).

**Figure 9.**
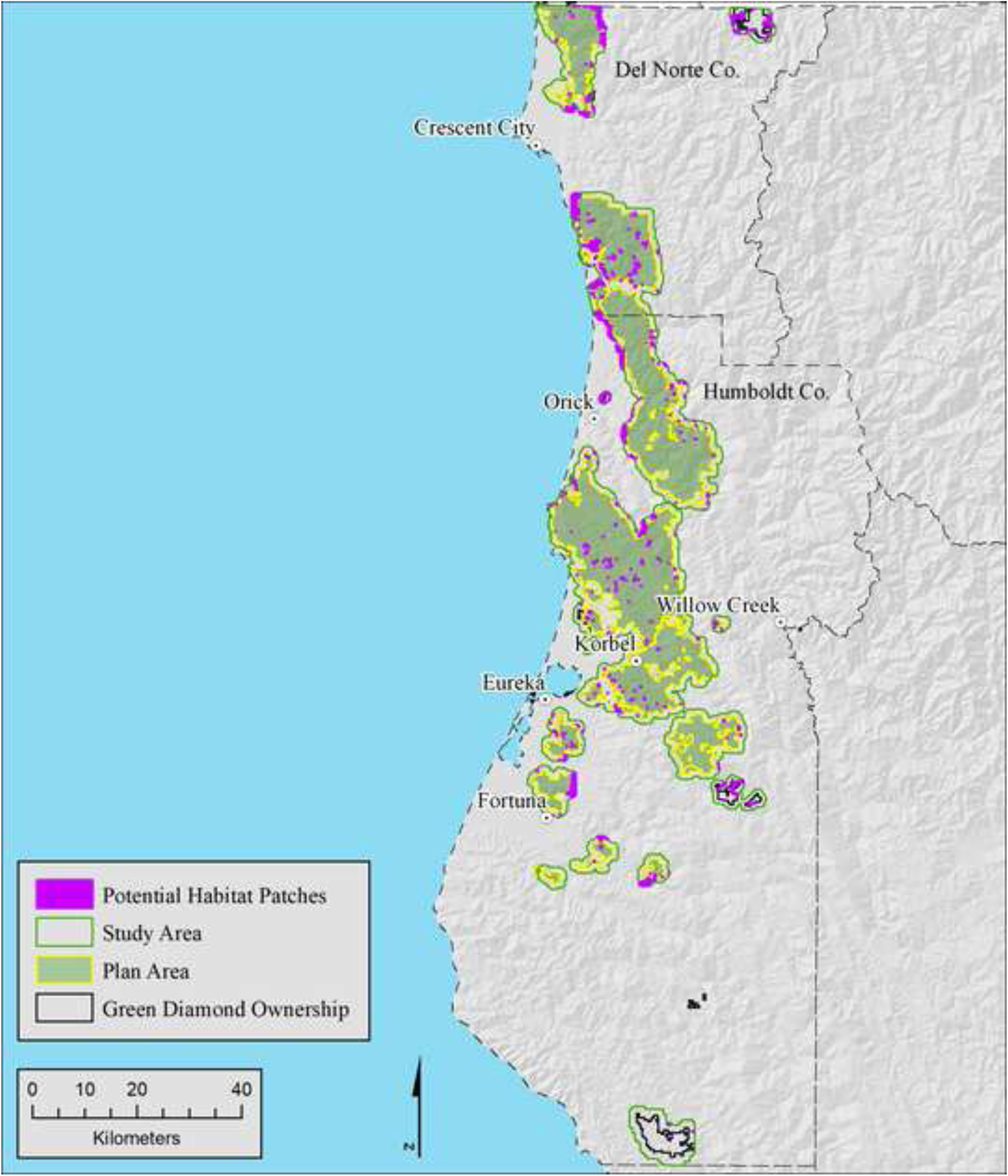
Potential habitat across the Study Area. Distribution of potential habitat patches within the Study Area. Patches were identified via a concave hull process applied to all TAO’s within a five-acre buffer (80.25m radius) of areas identified as low suitable or better (HSI ≥ 0.19) by the Maxent model.

**Table 3.** Summary of model delineated habitat. The values represent the model predicted area by suitability classes and occupancy status, rounded to the nearest whole hectare, based on the raw HSM raster. ‘Used’ patches are those with documented occupied behaviors, and within the Study Area include patches on Federal and State parks, Bureau of Land Management reserves, unharvested old growth private managed forests, partially harvested old growth forests, and younger (< 70 years old) second and third growth forests with residual old growth trees. ‘Available’ patches are areas with non-confirmed Occupancy (Occupied behaviors) status. Both ‘Used’ and ‘Available’ are delineated by methods described in the text.

|  | **Plan Area (Green Diamond only)** |  |  |  |  |  |
| --- | --- | --- | --- | --- | --- | --- |
|  | **Unsuitable** | **Marginal** | **Low** | **Medium** | **High** | **Plan Total** |
| Used | 16 | 86 | 70 | 96 | 24 | 292 |
| Available | 21 | 270 | 118 | 29 | 0 | 437 |
| Total | 37 | 356 | 188 | 125 | 24 | 729 |
|  | **Study Area (Green Diamond plus ½-mile off-property buffer)** |  |  |  |  |  |
|  | **Unsuitable** | **Marginal** | **Low** | **Medium** | **High** | **Study Total** |
| Used | 109 | 377 | 339 | 627 | 1,465 | 2917 |
| Available | 172 | 1,083 | 620 | 466 | 163 | 2504 |
| Total | 281 | 1,460 | 960 | 1092 | 1628 | 5,421 |

In the Plan and Study Areas, polygon patches we considered occupied comprised 89.9% and 99.5%, respectively, of the raster-based high habitat classification, 54.2% and 66.8% of the medium classification, and 30.7% and 28.9% of the low classification. Not surprisingly, these figures indicate that most of the better murrelet habitat occurred in the patches we considered occupied, and hence few high-quality patches exist outside our already identified ‘used’ patches. Small areas of low, medium, or high suitability were occasionally left outside of concave hull delineated patches because they lacked 50-meter and taller trees (Figure 4).

### Model Evaluation

Our final HSM model attained an AUC value of 0.99 (Table 4, Figure 10). Furthermore, AUC varied little over the ten bootstrap replicates evaluated by Maxent. AUC = 0.99 indicates a model with very high predictive ability and low probability of false negatives. We adopted the lower limit of the Low habitat class (i.e., HSI = 0.19) as the threshold above which we classified an area as murrelet habitat. With this threshold, our model achieved a sensitivity value of 0.89 indicating that nearly all ‘used’ locations had HSI values above the threshold and were predicted to be habitat by the model. Our model attained a Positive Predictive Value (PPV) of 0.49 indicating a significant number of ‘available’ locations (987) with HSI values above the threshold (0.19). Our model achieved a Cohen’s Kappa value of 0.63 and indicated “substantial” agreement between the predicted habitat areas and known ‘used’ locations if we adopt the guidelines of Landis and Koch (1977). Other authors consider Cohen’s Kappa = 0.63 to be “good” agreement (Fleiss, 1981; Luck, 2002).

**Figure 10.**
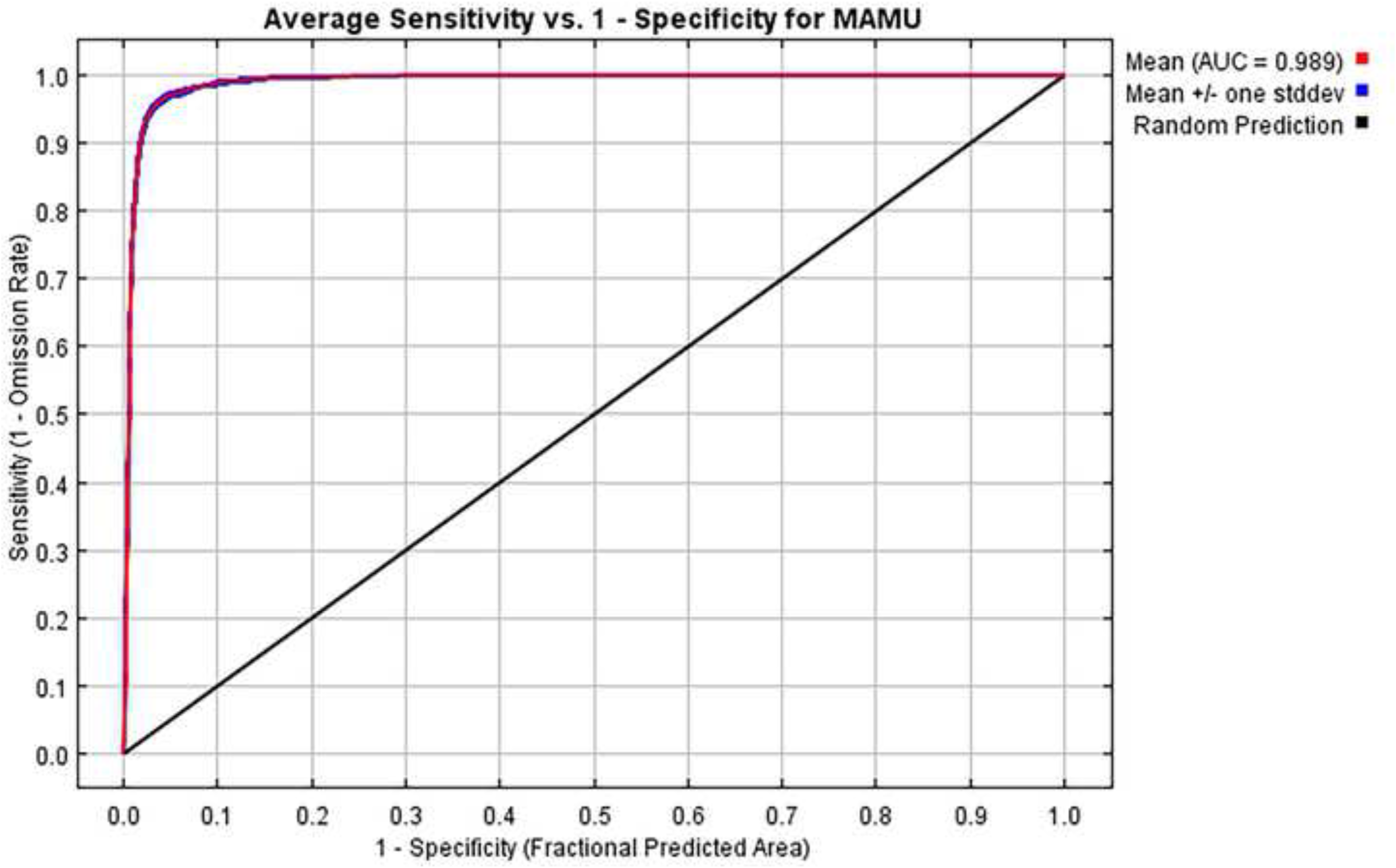
Receiver Operating Characteristic (ROC) curve. This ROC curve is an average of 10 replicate runs. The average training Area Under the ROC Curve (AUC) for the replicate runs is 0.989, and the average standard deviation is 0.001. The best model in our analysis predicting the environmental niche of marbled murrelets for the study area had an AUC value of 0.989 and exhibited very low deviation from the mean of the ten replicates over the full ROC curve. This value indicates a model with high predictive ability and low probability of false negatives.

**Table 4.**
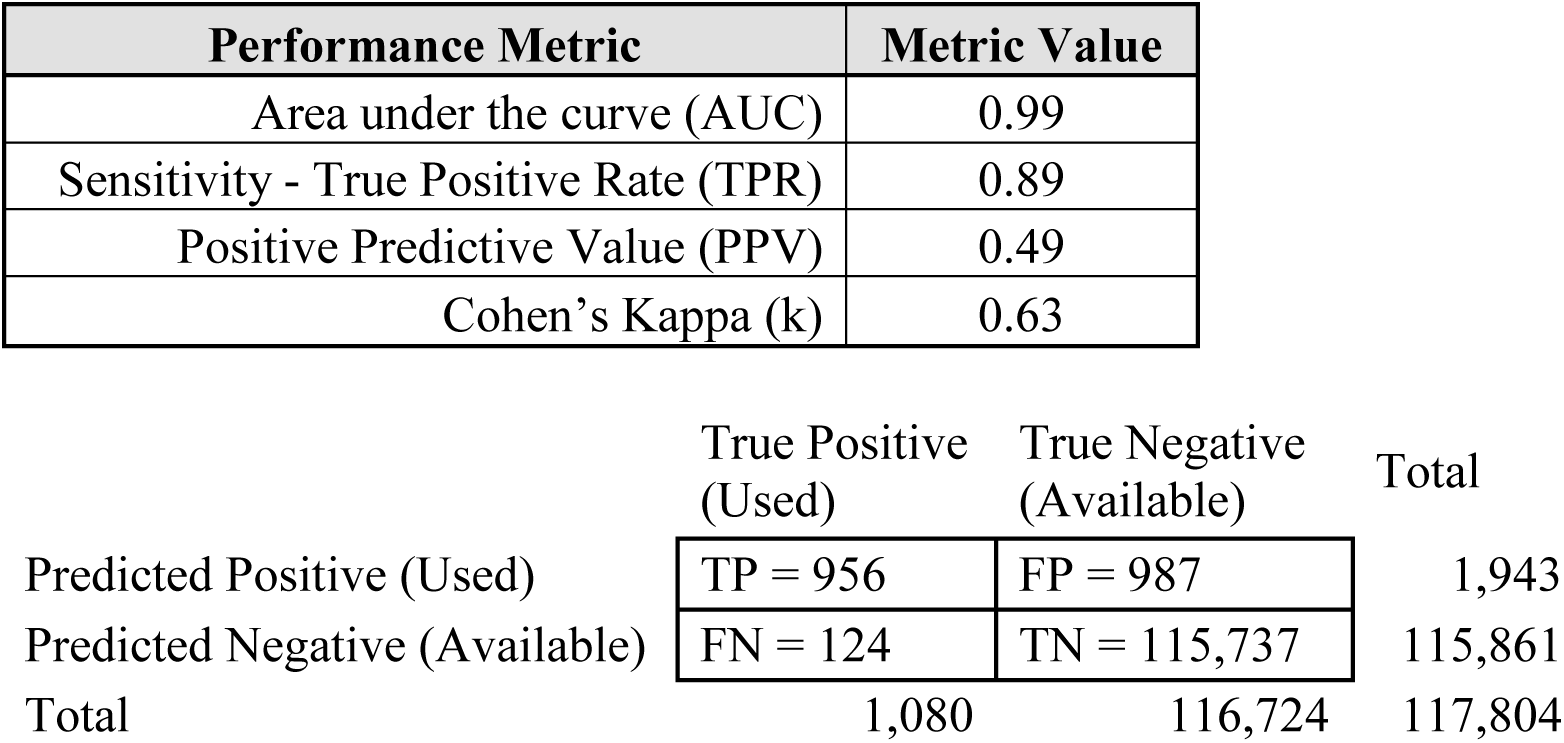
Model performance measures. Predictive performance measures for the final model of marbled murrelet habitat using training data and HSI threshold of 0.19. Sensitivity is the number of correctly predicted used locations divided by the total number of locations predicted to be “habitat” (i.e., TP / (TP + FN) = 956 / 1,080). Positive predictive value is number of correctly predicted used locations divided by total number of used locations (i.e., TP / (TP + FP) = 956 / (956 + 987). Cohen’s Kappa measures the agreement between known and predicted values and is defined as 2(TP * TN – FN * FP) / (TP + FP) * (FP + TN) + (TP + FN) * (FN + TN)) (i.e., 221,044,368 / 351,924,612).

| Performance Metric | Metric Value |
| --- | --- |
| Area under the curve (AUC) | 0.99 |
| Sensitivity - True Positive Rate (TPR) | 0.89 |
| Positive Predictive Value (PPV) | 0.49 |
| Cohen’s Kappa (k) | 0.63 |

|  | True Positive<br>(Used) | True Negative<br>(Available) | Total |
| --- | --- | --- | --- |
| Predicted Positive (Used) | TP = 956 | FP = 987 | 1,943 |
| Predicted Negative (Available) | FN = 124 | TN = 115,737 | 115,861 |
| Total | 1,080 | 116,724 | 117,804 |

## Discussion

Our ‘used’ habitat records on private timberland were derived from focused surveys conducted by trained individuals after input and guidance from regulatory agencies. We surveyed a wide and representative range of environmental conditions within the study area to establish used patches because these surveys were necessary to obtain timber harvest permits. Our BAS sample minimized sampling bias of background values because points came from a spatially balanced sample of environmental conditions within and outside of known occupied habitats.

Within the California range of marbled murrelets, only 26 nest trees have ever been located and only 1 fell on private timberland in our study area. These 26 nest trees were used in analyses conducted for the Northwest Forest Plan (Lorenz et al., 2021), and these authors supplemented their small sample size by including additional information from eggshell fragments (n=56), downed young (n=11), and an equivalent total number of occupied sites (n=93). Like Hamer and Nelson (1995) and Lorenz et al. (2021), we used several sources of information to determine habitat use. We sourced most information from inland audio-visual surveys conducted on private timberland prior to timber harvest plan approvals from 1992 to 2021. During this period, we conducted audio-visual surveys in all areas that the landowner, state, and federal agencies agreed could potentially be murrelet habitat. Hence, the surveyed areas covered all potential murrelet habitat based on the prevailing knowledge of habitat during the study period. In total, we surveyed 26 such areas and our model ended up classifying these areas as ranging from unsuitable to medium, with most classified as unsuitable or marginal.

We do not know the degree to which detection probability varies with the covariates influencing the relative probability of occupancy (Yackulic et al., 2013). However, on private timberlands we used data from surveys designed to clear areas for timber harvest and that were designed to detect murrelet presence with 95% or greater accuracy. Because of this, we discount the possibility that a large number of murrelets were not detected on private timberlands and that the rare non-detection depended on covariates affecting selection. We also discount the possibility that high densities of large trees reduce detection probability because the audio-visual surveys were specifically designed to detect murrelets in these situations.

We used a very large ‘available’ sample which has been noted to artificially improve AUC simply because it is partially dependent on the ratio of ‘used’ to ‘available’ points (Yackulic et al., 2013). Our large ‘available’ sample was necessary to adequately quantify the distribution of extremely patchy and sometimes rare environmental conditions relevant to marbled murrelets. To assess whether our large ‘available’ sample artificially inflated AUC, we refitted the final model after randomly reducing the ‘available’ sample from 117,804 to 11,828 records (10.04% of the original). The refitted final model yielded an AUC of 0.987 compared to the original model’s AUC = 0.989. Likewise, we simulated the effects of smaller ‘available’ samples on Cohen’s Kappa by successively halving the ‘available’ sample. Contrary to expectations, Kappa increased from 0.63 (117,804 ‘available’ points) to a maximum of 0.90 (7,295 ‘available’ points – close to the Maxent default of 10,000) then decreased to 0.72 (228 ‘available’ points). While variable, all simulated Kappa values suggest substantially better prediction than random classification. Generally, Kappa (κ) values less than 0.4 are considered poor, 0.4 < κ < 0.75 are considered good, and κ greater than or equal to 0.75 excellent (Cohen, 1960; Landis and Koch, 1977; Fleiss, 1981; Luck, 2002). Our reported value of 0.63 falls in the “good” range.

Our HSM scored very high to reasonably well on the predictive accuracy measures we computed. The principal issue impeding higher sensitivity scores is the existence of used locations with HSI values below the threshold of 0.19 (i.e., false negatives) affecting approximately 11% of our used sample. Two conditions resulted in false negatives. The foremost was due to forest heterogeneity within the occupied patches falling mostly on managed timberlands. Within the ‘used’ patches, internal and edge inclusions of low-quality habitat (sparse large trees) with HSI values below the threshold were present in approximately 8% of the used sample. A less important effect occurred near the abrupt edge between old trees and surrounding young forest. Model selection supported a 2-hectare (5-acre, 80.25-meter radius) moving window to calculate predictor variables resulting in gradual decreases in HSI values as the circular window moved out of presumed nesting patches and into adjacent younger forest (Figure 11). This decline near patch edges results in approximately 3% of our used locations near patch edges being associated with HSI values below the threshold.

**Figure 11.**
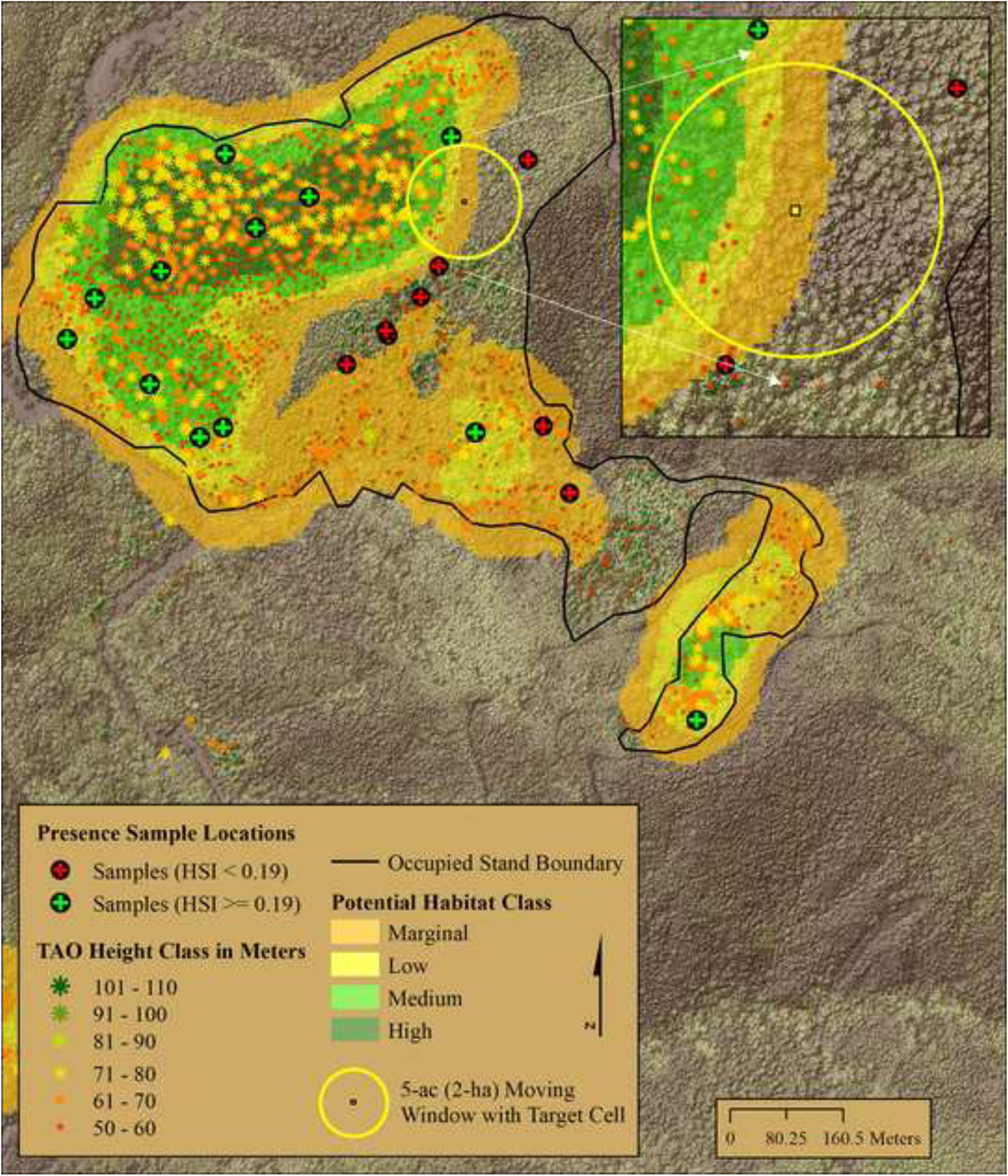
BAS training samples. An example of BAS sample locations used to train the Maxent model, TAOs ≥ 50 meters tall, potential habitat classifications, and 5-acre circular moving window with central target cell. As the 5-acre circular window moves from a core area containing higher densities of TAOs ≥ 50 meters to non-habitat area with low densities or no TAOs ≥ 50 meters outside the occupied stand, the modeled HSI values calculated for the target cell decrease as a result. This heterogenous distribution of TAOs ≥ 50 within occupied stands results in internal pockets and areas adjacent to the occupied stand edges with HSI values below the threshold value of 0.19 leading to perceived errors of omission (black circle/red +).

PPV is directly affected by the presence of predicted habitat outside of the ‘used’ or known occupied patches. Our relatively low PPV value is typical in ‘used’ - ‘available’ HSM predictive efforts where the extent of true occupancy is likely greater than known prior to modeling and when the sample of used locations is small relative to the study area.

Our correlative HSM model indicated that fine-scale, LiDAR-based, individual tree covariates were the best predictors of habitat suitability for marbled murrelets. We summarized height and density of our derived individual trees (TAOs) at two different scales, 1-acre, and 5-acre, using moving window methods that best represent existing known and potential habitat patches within the study area. During model selection, the 5-acre scale outperformed covariates compiled at the 1-acre scale and no climatic or biographic/geographic themed variable performed well enough to be included in the final phase. We conclude that the 5-acre scale is a better predictor of marbled murrelet habitat within managed landscapes where environmental covariates are patchily distributed and more dispersed than in large homogeneous old growth stands. We hypothesize that low variation in the climatic, biographic, and geographic variables at a study area scale rendered these covariates ineffective as predictors. We suspect habitat selection could be associated with natural geographic factors such as topographic position and distance to coast (which is positively correlated with elevation) as found in other studies (Hagar et al., 2014; Clyde, 2017); however, the entire relevant range of geographic and topographic conditions was not well represented in our study and hence the variable selection procedure did not include these factors in the model.

In Oregon, Hagar et al. (2014) used a suite of Gradient Nearest Neighbor (GNN) covariates and LiDAR derived covariates to develop a logistic regression model predicting relative probability of marbled murrelet occupancy in similar aged forest stands that were all considered potential habitat based on age (> 80 years) and structural characteristics. They found that the model with five LiDAR derived variables along with distance to the coast performed better than the model with GNN derived variables. The LiDAR variables they used were better predictors of habitat for marbled murrelets at stand and watershed scales when applied to independent data sets, although overall performance was low on the independent data set. They interpreted the LiDAR-derived variables in their model as cover in the upper portion of the canopy, maximum tree height, maximum height of the bottom of the canopy, variation in vertical stratification of the canopy, and distribution of variation across canopy height intervals. These findings are consistent with studies that found multi-layered old forests provide the components necessary to support nesting by marbled murrelets (Hamer and Nelson, 1995; Nelson and Wilson, 2002; Raphael et al., 2011). Clyde (2017) assessed the use of individual tree objects, modeled from LiDAR to calculate the local Moran’s I spatial statistic (LMI), to identify the presence of vertically isolated trees and or clusters of tall trees. While the LMI metric yielded low predictive power, canopy height was shown to be important for the highest ranked areas.

Finally, our study advances understanding of marbled murrelet habitat use by incorporating fine scale LiDAR derived individual tree metrics (height, height diversity and density) directly into predictive covariates. All three LiDAR variables included in our HSM are more direct measures of small-scale marbled murrelet habitat requirements than indirect area-based point cloud metrics derived at various raster scales and resolutions. However, because our data contains information on tall trees ≥ 50 meters, field assessments of the presence, quantity, and quality of platforms may facilitate identification of habitat because young tall trees might not provide suitable platforms for nesting (Sillett et al., 2018). In the future, LiDAR point cloud data may provide platform information on individual trees that will inform future marbled murrelet HSMs.

## Acknowledgements

We thank Green Diamond Resource Company for supporting this research and specifically Galen Schuler for his invaluable input and encouragement during this endeavor. In addition, we thank Jake Verschuyl, PhD, Richard Golightly, PhD, and Tim Bean, PhD for insightful review of the initial draft of the document.

